# Sequential transcriptional programs underpin activation of quiescent hippocampal stem cells

**DOI:** 10.1101/2024.10.29.620545

**Authors:** Piero Rigo, Sara Ahmed-de-Prado, Rebecca L. Johnston, Chandra Choudhury, François Guillemot, Lachlan Harris

## Abstract

Postnatal neural stem cells are primarily quiescent, which is a cellular state that exists as a continuum from deep to shallow quiescence. The molecular changes that occur along this continuum are beginning to be understood but the transcription factor network governing these changes has not been defined. We show that these transitions are regulated by sequential transcription factor programs. Single-cell transcriptomic analyses of mice with loss- or gain-of-function of the essential activation factor *Ascl1*, reveal that Ascl1 promotes the activation of hippocampal neural stem cells by driving these cells out of deep quiescence, despite its low protein expression. Subsequently, during the transition from deep to shallow quiescence, *Ascl1* induces the expression of *Mycn*, which drives progression through shallow states of quiescence towards an active state. Together, these results define the required sequence of transcription factors during hippocampal neural stem cell activation.

## INTRODUCTION

Cellular quiescence is not a single state but a spectrum. Stem cells that are in shallow quiescence, for example in a G_alert_ state, have short activation times, whereas deeply quiescent cells require prolonged growth factor signalling to activate^1^. The different states of quiescence have functional significance. For example, tissue injury pushes muscle stem cells and haemotopioetic stem cells in uninjured areas of the body into a shallow quiescent state, thereby priming these cells for regenerative activity^2^. In contrast, fasting pushes muscle stem cells into deep quiescence, decreasing regenerative capacity but increasing the resilience of the cells to stress^3^.

Likewise, in the central nervous system, quiescent neural stem cells (NSCs) exhibit different depths of quiescence. Injury pushes quiescent NSCs in the subventricular zone into a ‘primed’ state of shallow quiescence^4^. NSCs that return to quiescence from a proliferating state enter a shallow ‘resting’ state of quiescence, which increases the probability that these cells will re-enter the cell-cycle in the short term compared to cells that have not divided recently^5,6^. Finally, NSCs also progress into deeper states of quiescence the longer they remain quiescent, which reduces the activation rate of NSCs in aging animals^6–8^.

Despite the functional significance of these different states, how adult NSCs convert between states of deep and shallow quiescence is poorly defined. Here, we reveal that in adult mouse NSCs, activation from deep quiescence and shallow quiescence is governed by distinct and sequential molecular programs. Progression from deep quiescence to shallow quiescence is promoted by the *Ascl1* transcription factor. ASCL1 induces the expression of *Mycn* to push cells through shallow states of quiescence into an active state. These defined transitions may explain how populations of hippocampal NSCs are maintained in distinct states of quiescence.

## RESULTS

### Ascl1 deletion impairs progression from deep states of quiescence

Expression of the transcription factor ASCL1 is essential for the activation of adult hippocampal NSCs^9^. ASCL1 must also be degraded for active NSCs to return to a quiescent state^10^. While analysis of ASCL1 function has focused so far on its role in the active NSCs, there is clear evidence that ASCL1 is also expressed in quiescent NSCs, albeit at lower levels, raising the possibility that it has a role in this population^6,11^. To determine how ASCL1 might function during quiescence, we performed single-cell RNA-sequencing (scRNA-seq) of hippocampal NSCs after acute deletion of the *Ascl1* gene.

We administered tamoxifen to Ascl1*fl/fl*; Glast-creERT2; Rosa-YFP mice (*Ascl1* cKO) at 1- month of age, deleting the *Ascl1* gene from NSCs and parenchymal astrocytes, while irreversibly labelling recombined cells with yellow fluorescent protein (YFP)^9,12,13^. Twelve-days after tamoxifen administration, we disassociated the dentate gyrus of *Ascl1* cKO mice, as well as control animals that lacked the conditional *Ascl1* allele. We sorted the recombined YFP+ *Ascl1* cKO and control cells by flow cytometry and performed scRNA-seq with the 10x Genomics platform (Figure 1A, Table S1). In total, 12,548 cells from *Ascl1* cKO mice and 9,189 cells from controls, across three independent experiments passed quality control. We first inspected our data by isolating the three main neurogenic cell clusters (Figure 1B). These clusters comprised quiescent NSCs, proliferating cells (both active NSCs and intermediate progenitor cells, which cluster together due to dominance of cell-cycle gene expression) and neuroblasts, which we demarcated according to the expression of canonical markers *Hopx*^14^, *Eomes/MKi67* and *Dcx*^15^ respectively, as previously described^6^. Consistent with the established role for ASCL1 in active NSCs^9^, loss of ASCL1 led to an almost complete absence of actively dividing cells in *Ascl1* cKO mice (Figure 1C). In our scRNA-seq data of *Ascl1* cKO mice, only 8 cells were actively proliferating (0.55%) compared to 20% of control cells (Figure 1D). We have previously reported that this block in proliferation leads to an accumulation of NSCs in a quiescent state because of a lack of division-coupled depletion^9,16^. Indeed, in our scRNA-seq data 96.2% of cells were quiescent NSCs compared to 25.3% of control cells (Figure 1D). Finally, the block in proliferation also leads to a histological loss of immature neuronal progeny^9^ and in our scRNA-seq data just 3.27% of cells in *Ascl1* cKO mice were neuroblasts compared to 54.8% of controls cells (Figure 1D). The small number of aberrant proliferating cells and neuroblasts in our scRNA-seq data from *Ascl1* cKO mice was likely due to the incomplete recombination events^9^, as we detected *Ascl1* mRNA expression in the majority of remaining proliferating cells in the *Ascl1* cKO dataset. Our scRNA-seq data therefore faithfully recapitulates the histological phenotype of *Ascl1* cKO animals^9^, allowing us to use this sequencing tool to determine how *Ascl1* loss affects the underlying biology of the quiescent NSC population.

**Figure 1:**
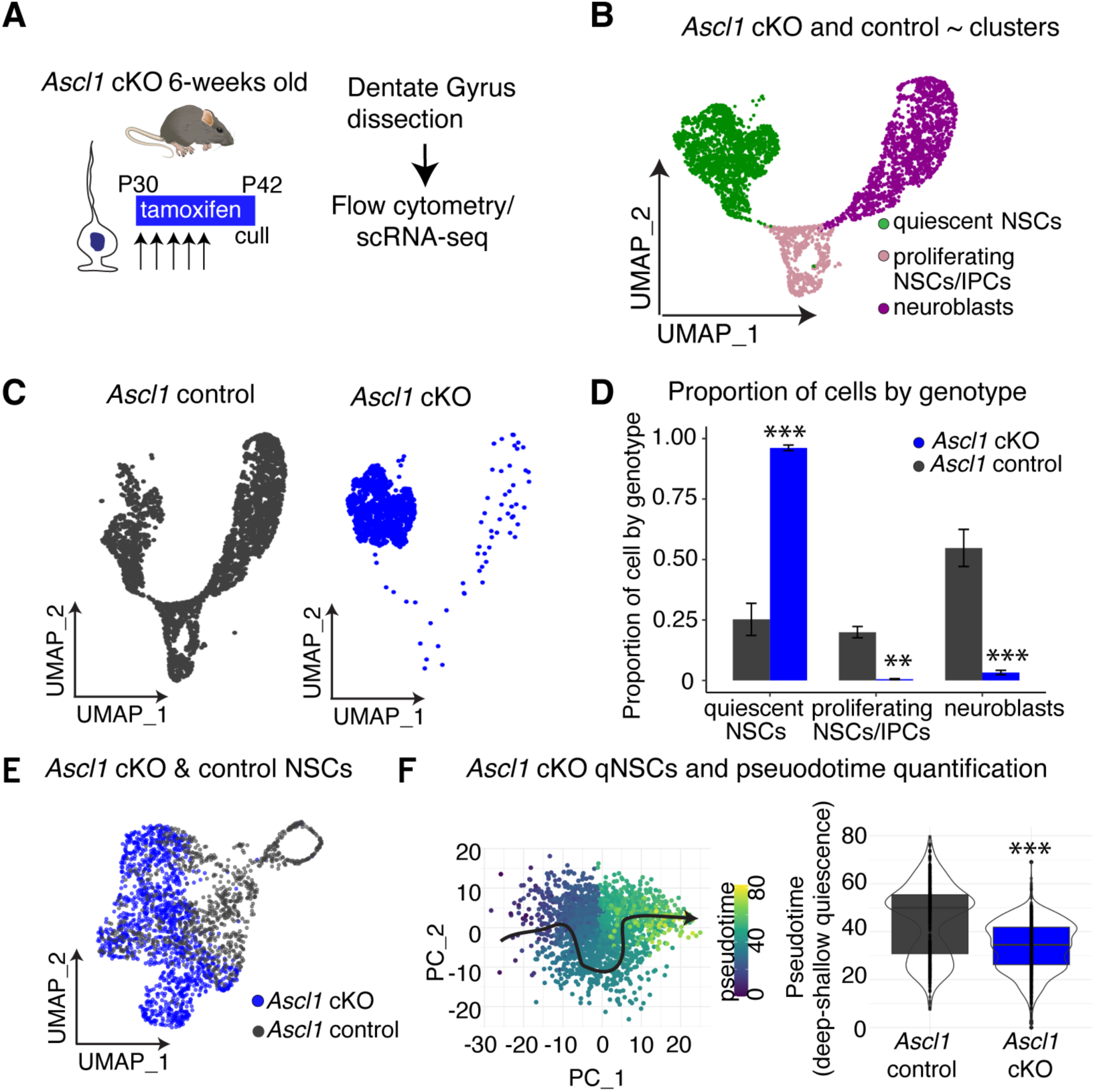
*Ascl1* loss reduces NSC activation by impeding progression from deep states of quiescence. (A) To assess how loss of ASCL1 affects quiescence, mouse hippocampal dentate gyrus was dissected, and genetically recombined cells (Glast-creERT2; RosaYFP) lacking *Ascl1* (*Ascl1* cKO) and controls were processed for scRNA-seq. (B) UMAP of single cell transcriptomes from *Ascl1* cKO and control mice showing quiescent NSCs, proliferating cells (NSCs and intermediate progenitors) and neuroblasts. (C) UMAP of single cell transcriptomes from *Ascl1* cKO and control mice split by genotype. (D) Quantification of the proportion of quiescent NSCs, proliferating cells and neuroblasts in *Ascl1* cKO and control mice. (E) UMAP of NSCs from *Ascl1* cKO and control mice. (F) Pseudotime analysis of quiescent NSCs from *Ascl1* cKO and control mice. Left: Analysis visualised using first two principal components, with cells coloured by pseudotime and cell ordering indicated by black arrow. Right: Distribution of pseudotime values per genotype. Statistics: Kolmogorov-Smirnov test in (F), multiple comparison t-test in (D) reporting Holm-Sidak corrected *P*-value. \*\**P* < 0.01, \*\*\**P* < 0.001. Abbreviations: intermediate progenitor cell (IPC).

To examine the effect of *Ascl1* deletion on the quiescent stem cell population, we re-clustered the scRNA-seq data to only include NSCs (*Hopx*-high; *S100b*-low) (Figure 1E) and then only quiescent NSCs, by removing proliferating cells (Figure 1F). We arranged the quiescent NSCs along a pseudotime axis from deep to shallow quiescence using the trajectory inference tool Slingshot^17^ (Figure 1F). Loss of ASCL1 led to a pronounced phenotype in quiescent NSCs, where much of the NSC population shifted to extremely deep states of quiescence. Specifically, almost all quiescent NSCs from *Ascl1* cKO mice occupied the deepest 50% of pseudotime positions from control mice (Figure 1F). These data demonstrate that loss of ASCL1 in adult NSCs causes profound transcriptional changes to all NSCs, including quiescent NSCs where it is expressed at low levels, ultimately moving the population into a deeper quiescent state. We next tested whether, conversely, increasing the level of ASCL1 protein might shift deeply quiescent NSCs to a shallow state.

### Ascl1 gain-of-function promotes progression from deep states of quiescence

To perform an ASCL1 gain-of-function analysis, we repeated the scRNA-seq experimental design, except this time deleting the gene encoding the ubiquitin-ligase HUWE1 (Figure 2A). The protein HUWE1 marks ASCL1 for proteasomal degradation. Our previous work has demonstrated that ASCL1 is the major substrate of HUWE1 in adult hippocampal NSCs and thus its deletion leads to elevated levels of ASCL1 protein in active NSCs^10^ and in quiescent NSCs^6^. The scRNA-seq analysis of dentate gyrus cells isolated from two independent experiments of *Huwe1* cKO mice (6,784 cells) and controls (5,187 cells) recapitulated our previous immunolabeling characterisation of this strain^10^. Specifically, a much larger percentage of cells from the neurogenic lineage were proliferating in *Huwe1* cKO mice (44.5% of cells) compared to control cells (13.6% of cells) (*P* = 0.08; Figure 2B-D). Furthermore, as we have previously shown, *Huwe1* cKO neuroblasts fail to exit the cell-cycle and differentiate, resulting in apoptosis. This phenotype was manifested in the scRNA-seq analysis as a reduction in the proportion of neuroblasts (*P* = 0.08; Figure 2B-D). We then re-clustered the scRNA-seq data to only include NSCs (Figure 2E) and then only quiescent cells (Figure 2F) and organised them into a pseudotime trajectory. We found that the increase in proliferation in *Huwe1* cKO mice was due to a marked shift in the stem cell population so that more mutant cells fell within the shallow end of the quiescence-to-activity trajectory and were likewise depleted from deep states of quiescence (Figure 2F). Together, the pseudotime trajectories of *Ascl1* loss- and gain-of-function mice reveal a surprisingly early role for ASCL1 in the activation process. Despite the low levels of ASCL1 protein in deeply quiescent NSCs, ASCL1 is required to promote the progression of these cells from deep to shallow quiescence. These results help to explain our previous observations that NSCs progress into deeper states of quiescence during aging and that this was tightly correlated with decreasing ASCL1 protein levels in quiescent hippocampal NSCs in older mice^6^.

**Figure 2:**
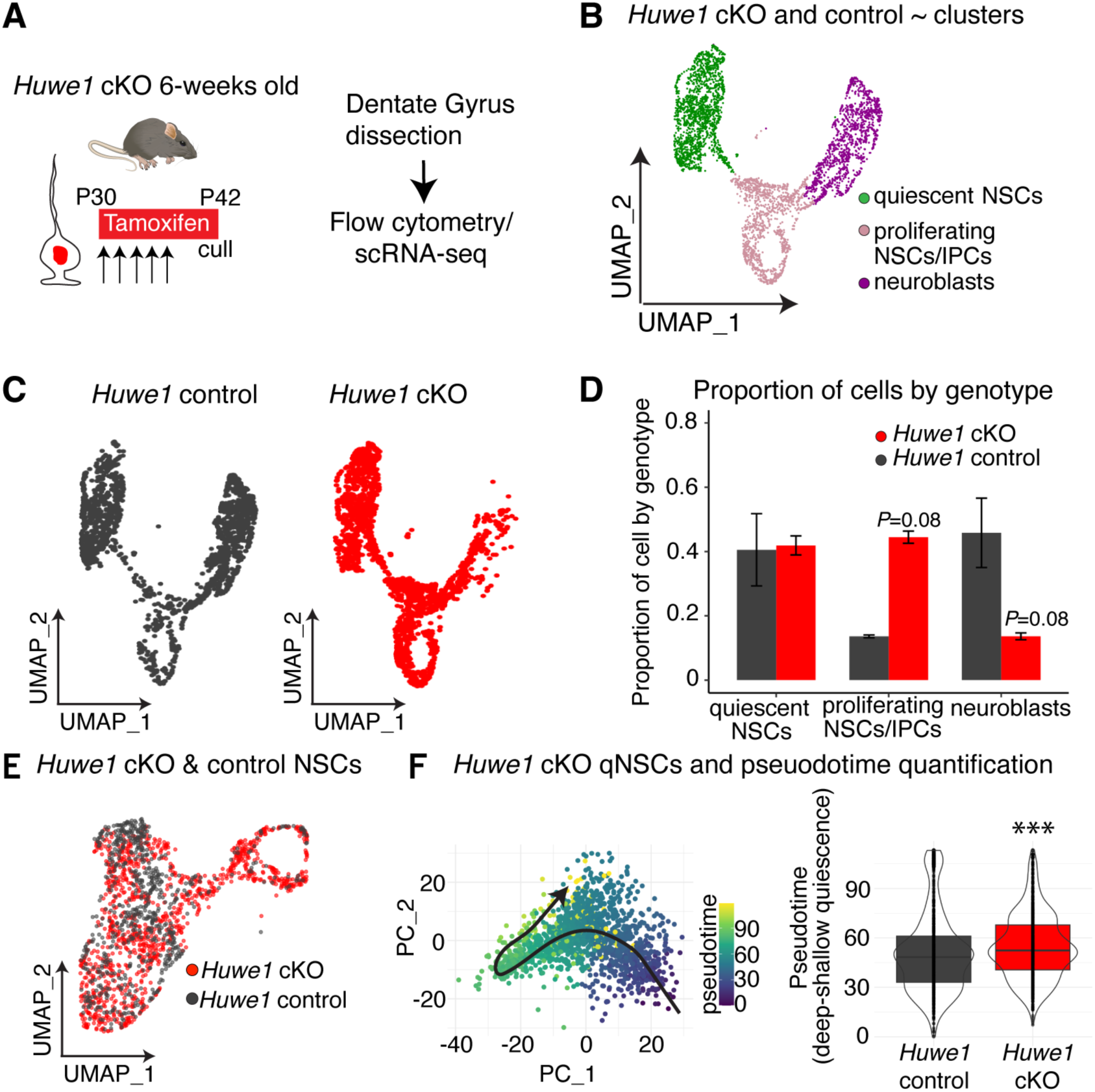
Ascl1 gain-of-function increases NSC activation by promoting progression from deep states of quiescence. (A) To assess how loss of HUWE1 affects quiescence, mouse hippocampal dentate gyrus was dissected, and genetically recombined cells (Glast-creERT2; RosaYFP) lacking *Huwe1* (*Huwe1* cKO) and controls were processed for scRNA-seq. (B) UMAP of single cell transcriptomes from *Huwe1* cKO and control mice showing quiescent NSCs, proliferating cells (NSCs and intermediate progenitors) and neuroblasts. (C) UMAP of single cell transcriptomes from *Huwe1* cKO and control mice coloured by genotype. (D) Quantification of the proportion of quiescent NSCs, proliferating cells and neuroblasts in *Huwe1* cKO and control mice. (E) UMAP of NSCs from *Huwe1* cKO and control mice. (F) Pseudotime analysis of quiescent NSCs from *Huwe1* cKO and control mice. Left: Analysis visualised using first two principal components, with cells coloured by pseudotime and cell ordering indicated by black arrow. Right: Distribution of pseudotime values per genotype. Statistics: Kolmogorov-Smirnov test in (F), multiple comparison t-test in (D) reporting Holm-Sidak corrected *P*-value. \*\*\**P* < 0.001. Abbreviations: intermediate progenitor cell (IPC).

### MYCN promotes progression from shallow states of quiescence

We next wondered whether the subsequent progression of adult NSCs through shallow quiescence and into an active state might be associated with the activity of transcription factor(s) other than ASCL1. NSCs in shallow quiescence such as primed and resting cells have higher expression of genes associated with mRNA transport and protein translation^4,6,18^. These pathways have been shown to be upregulated by *Myc* transcription factors in other quiescence-associated contexts, such as during diapause in preimplantation embryos and in dormant haemopoietic stem cells^19,20^. This led us to hypothesise that the *Myc* gene family might similarly increase the expression of these pathways in quiescent NSCs to promote the activation of these cells out of shallow quiescence.

We first individually deleted the three *Myc* gene family members from adult stem cells and parenchymal astrocytes, by crossing conditional *Myc*^21^, *Mycn*^22^ and *Mycl* mice^23^ to *Glast-creERT2* animals. For each strain, we administered tamoxifen to 2-month-old animals, culling them 30-days later (Figure S1). Potential genetic redundancy between *Myc* family members was explored by generating a triple cKO strain (Figure S2)^24^. We stained brain slices for the immature neuronal marker DCX, as a preliminary screen for underlying defects in NSC activation, which, all else being equal, would be expected to culminate in reduced levels of adult neurogenesis. Of the three family members, *Mycn* had the highest mRNA expression in cells of the neurogenic lineage (Figure S1). Consistent with this, deletion of *Mycn* from adult NSCs, though not that of *Myc* or *Mycl*, led to a substantial reduction in DCX-positive cells compared to control mice (Figure S1). Conditional deletion of all three *Myc* genes did not exacerbate the phenotype beyond that of the single *Mycn* gene deletion (Figure S2). Therefore, *Mycn*, but not the other *Myc* genes, has an important role in promoting adult neurogenesis in the mouse hippocampus.

We next examined whether the phenotype of *Mycn* cKO was brought about by an underlying impairment in the NSC activation process. The expression of MYCN protein supported a possible role in regulating NSC activation, with 20% of quiescent NSCs (radial GFAP+SOX2+ cells) expressing the protein in *Mycn* control mice at 6-weeks of age (Figure S3A, B), which largely reflected the mRNA expression in a subset of NSCs in shallow quiescence (Fig S1G). Quantification of NSCs revealed that deletion of *Mycn* reduced the proportion of adult hippocampal NSCs that were proliferating by 80% (Figure S3C, D) but did not affect overall NSC number (Figure S3E). To define how Mycn loss leads to decreased proportion of proliferating hippocampal NSCs, we performed scRNA-seq of *Mycn* cKO mice and controls, in a design that mirrored that of *Ascl1* and *Huwe1* cKO experiments (Figure 3A). In total, across two independent experimental replicates, we sequenced 9,187 *Mycn* cKO cells and 6,845 control cells. Consistent with our immunolabelling data (Figure S2C; Figure S3D), an increased percentage of cells from *Mycn* cKO mice were quiescent NSCs compared to controls (62.3% versus 31.4%), and there were also fewer neuroblasts (22.5% versus 50.4%) (Figure 3B-D). To test how MYCN might affect NSC activation we then re-clustered the scRNA-seq data to only include NSCs (Figure 3E) and then only quiescent cells and organised them into a pseudotime trajectory (Figure 3F). Unlike NSCs from *Ascl1* cKO mice, which accumulated in deep states of quiescence, the *Mycn* cKO cells entirely overlapped with control cells. However, *Mycn* cKO were not phenotypically normal, instead they demonstrated a distinct phenotype, whereupon they accumulated approximately halfway along the pseudotime trajectory (Figure 3F). These data suggest that while both *Ascl1* and *Mycn* deletion leads to a loss or decrease in stem cell activation, these phenotypes arise, at least in part, through different cellular mechanisms. *Ascl1* loss principally hinders the ability of cells to leave deep quiescence, whereas loss of *Mycn* impedes progression through shallow quiescence. Consistent with this, using tradeSeq^25,26^ we found that the number of differentially expressed genes between Ascl1 *cKO* mice and controls were highest during early phases of pseudotime (deep quiescence, defined as cells positioned prior to the median pseudotime position of *Ascl1* control cells), while the number of differentially expressed genes in *Mycn* cKO mice and controls was highest in shallow quiescence (defined as cells positioned after the median pseudotime position of *Mycn* control cells) (Figure S4B, E).

**Figure 3:**
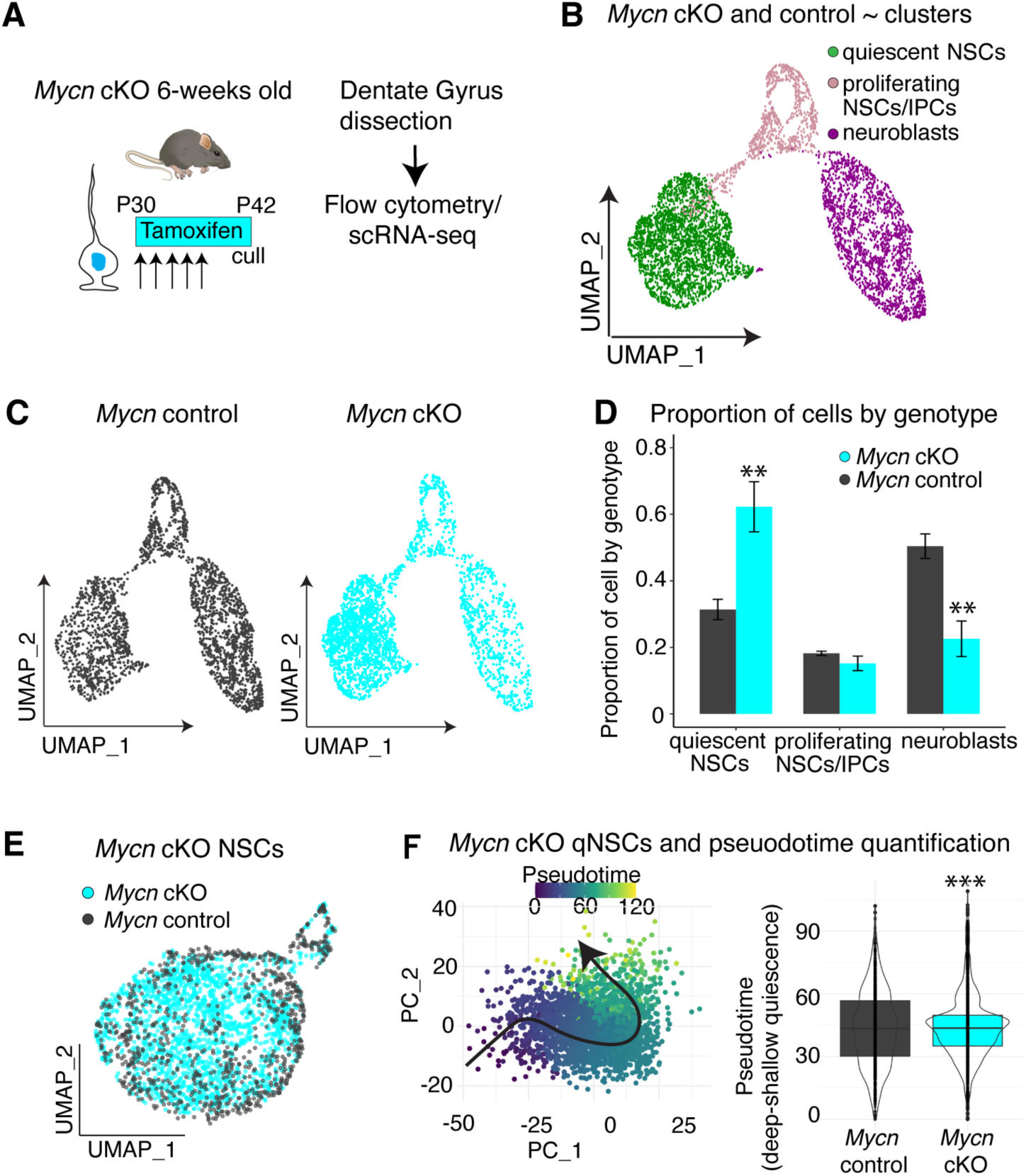
*Mycn* loss reduces NSC activation by impeding progression from shallow states of quiescence. (A) To assess how loss of MYCN affects NSC quiescence, mouse hippocampal dentate gyrus was dissected, and Nestin::GFP positive cells from *Mycn* cKO and controls were processed for scRNA-seq. (B) UMAP of single cell transcriptomes from *Mycn* cKO and control mice showing quiescent NSCs, proliferating cells (NSCs and intermediate progenitors) and neuroblasts. (C) UMAP of single cell transcriptomes from *Mycn* cKO and control split by genotype. (D) Quantification of quiescent NSCs, proliferating cells (NSCs and intermediate progenitors) and neuroblasts in *Mycn* cKO and control mice. (E) UMAP of NSCs from *Mycn* cKO and control mice. Abbreviations: intermediate progenitor cell (IPC). (F) Pseudotime analysis of quiescent NSCs from *Mycn* cKO and control mice. Left: Analysis visualised using first two principal components, with cells coloured by pseudotime and cell ordering indicated by black arrow. Right: Distribution of pseudotime values per genotype. Statistics: Kolmogorov-Smirnov test in (F), multiple comparison t-test in (D) reporting Holm-Sidak corrected *P*-value. \*\**P* < 0.01, \*\*\**P* < 0.001.

### Sequential Ascl1-Mycn programs govern NSC activation

The roles of ASCL1 and MYCN in promoting progression towards an active state from deep and shallow states of quiescence respectively, suggested that ASCL1 might induce the expression of MYCN during the activation process. Consistent with this, *Ascl1* mRNA levels increase dramatically prior to the induction of *Mycn* mRNA (Figure 4A, B). To explore this relationship further, we performed differential expression analysis on NSCs from *Ascl1* cKO mice and controls using a pseudo-bulk approach (Table S2). In NSCs from *Ascl1* cKO mice, *Mycn* was the second most strongly downregulated transcription factor in the dataset (Figure 4C, D; Table S2). We verified MYCN was also downregulated at a protein level in quiescent NSCs from *Ascl1 cKO* mice compared to controls (Figure 4E, F). Similarly, we performed differential expression analysis of NSCs from *Huwe1* cKO mice and controls (Table S3). In this analysis, loss of *Huwe1* led to a strong upregulation of *Mycn* (Figure 4G, H). Furthermore, we analysed a previously published ChIP-sequencing dataset of ASCL1 binding in hippocampal NSC cultures and found an ASCL1 peak overlapping with active enhancer marks P300 and H3K27ac in an intronic region of the *Mycn* gene (Figure 4I)^9^. Together, these data demonstrate that ASCL1 promotes the expression of *Mycn* during hippocampal NSC activation, likely through direct transcriptional activation.

**Figure 4:**
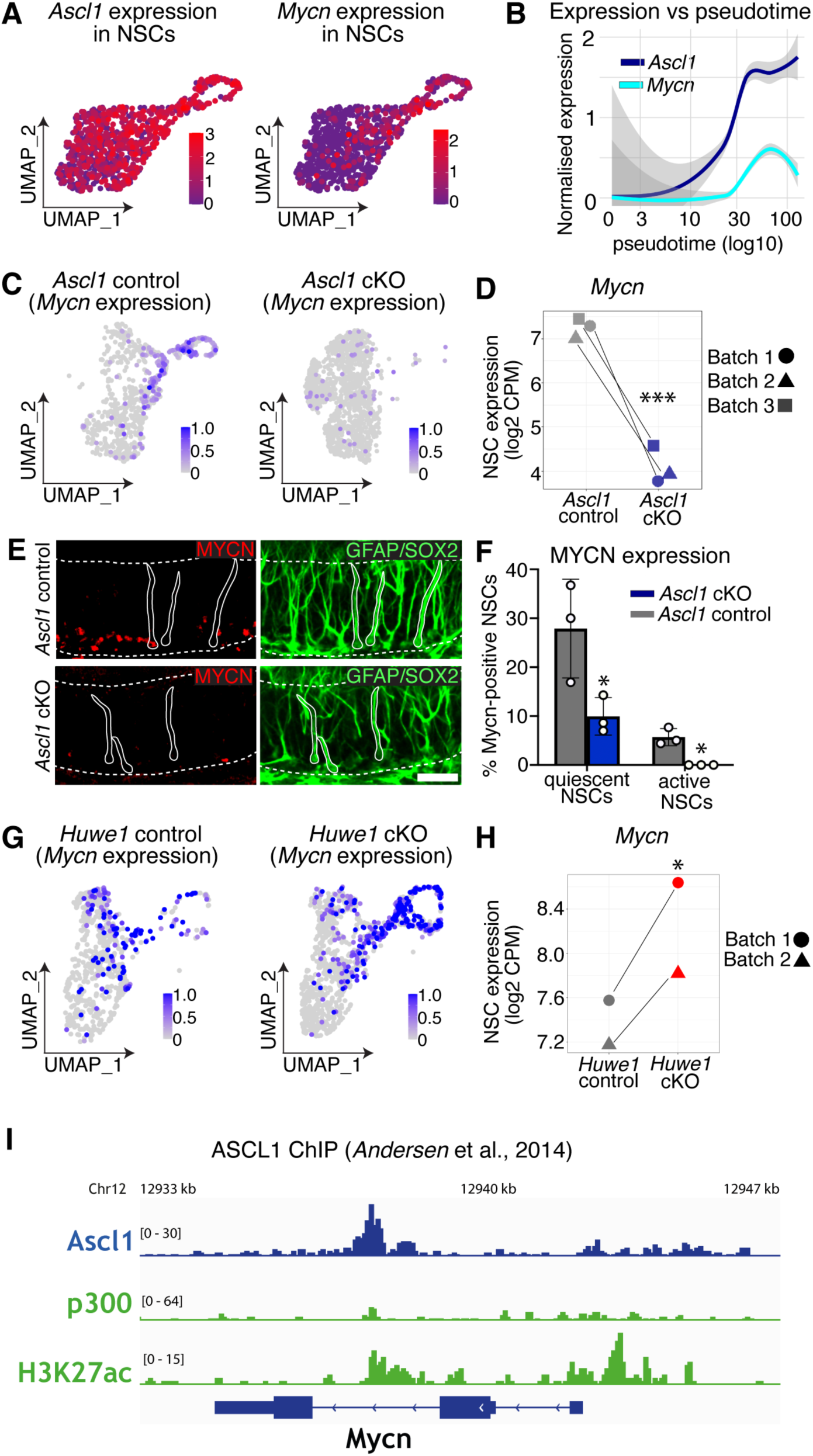
A sequential *Ascl1*-*Mycn* program drives NSC activation. (A) UMAP showing *Ascl1* and *Mycn* expression in wildtype NSCs. Data from *Ascl1* control mice in Figure 1. (B) *Ascl1* and *Mycn* expression in wildtype NSCs along pseudotime from deep quiescence to an active state. Data from *Ascl1* control mice in Figure 1. (C) UMAP plot showing *Mycn* expression in *Ascl1* cKO and control NSCs. (D) Scatterplot of *Mycn* expression in *Ascl1* cKO and control NSCs. Batch indicates experiment number. (E) MYCN protein is detected in fewer NSCs in *Ascl1* cKO NSCs than in *Ascl1* control NSCs. (F) Quantification of the proportion of NSCs that are quiescent and positive for MYCN (MYCN+KI67–) and the proportion that are active and positive for MYCN (MYCN+KI67+) in *Ascl1 cKO* and control mice. (G) UMAP plot showing *Mycn* expression in *Huwe1* cKO and control NSCs. (H) Scatterplot of *Mycn* expression in *Huwe1* cKO and control NSCs. Batch indicates experiment number. (I) ASCL1 ChIP data at *Mycn* locus in adult hippocampus derived NSCs. Dataset from Andersen et al., 2014^9^ and Martynoga et al., 2013^35^. Statistics: Pseudobulk differential expression in (D, H) reporting FDR-corrected *P* value; t test in (F). \**P <* 0.05, \*\*\**P* < 0.001. Scale bar (in E) = 30 µm.

We next sought to compare the transcriptional programs controlled by Ascl1 during deep quiescence and Mycn during shallow quiescence. To do this we performed differential expression analysis between cKO and control only on the quiescent NSC fraction from both the *Ascl1* datasets (Table S2, 158 differentially expressed genes) and *Mycn* datasets (Table S4, 92 differentially expressed genes). Gene ontology (GO) enrichment analysis of biological process terms (Table S2) revealed that loss of *Ascl1* led to enrichment of the *notch signalling pathway* (e.g., downregulation of genes *Dll1*, *Notch1/*2, *Hes5*) and *positive regulation of* C*yclin-dependent protein serine/threonine kinase activity* (e.g., downregulation of *Ccnd1/2* and *Egfr* genes). Loss of *Ascl1* also led to enrichment of *regulation of BMP signalling pathway*, including decreased expression of BMP antagonist *Nbl1* and altered expression of genes associated with *Cell-substrate adhesion* and *Mitochondrial membrane potential* (*Ndufc2, Mfn1*). In contrast, loss of *Mycn* resulted led to enriched GO terms relating to biosynthesis, specifically GO terms such as *cytoplasmic translation* (e.g., downregulation of *Rpl29, Rps12, Rps8* genes) and *regulation of transcription from RNA polymerase II promoter in response to stress* (e.g., *Egr1*, *Klf2, Pppr1r15a*) and (Table S4). No ribosomal genes or mRNA processing pathways were enriched in the *Ascl1* dataset (Table S2). These analyses demonstrate a clear separation of roles where ASCL1 promotes activation from deep quiescence by modulating the activity of key signalling pathways (e.g., cell adhesion and *BMP* signalling and metabolic modulation through mitochondria membrane potential)^27–29^, whereas MYCN promotes activation from shallow quiescence by increasing transcription and translation^18^.

## DISCUSSION

Quiescence is not a uniform cellular state; rather, it exists as a continuous spectrum of varying degrees of depth. Transitions between deep and shallow states of quiescence have been associated with the degree of lysosomal activity and autophagic flux^30^. However, outside of this, relatively little is known about the ordering of events during these transitions, including in the adult nervous system. Using scRNA-seq we have demonstrated that these transitions can be separated into distinct phases in quiescent hippocampal NSCs, where sequential activation of ASCL1 and MYCN transcription factors lead to NSC activation.

Prior work has focussed on the role of ASCL1 in active NSCs^9,10^, whereas our analysis has demonstrated a role for ASCL1 in quiescent NSCs. Specifically, our data demonstrates that ASCL1 is required to promote the progression of hippocampal NSCs from deep states to shallow states of quiescence. We have previously observed ASCL1 levels decline during aging and this is correlated with an accumulation of hippocampal NSCs in deep states of quiescence^6^. Our new data further strengthens the argument that this is a causal relationship. The role for ASCL1 in promoting activation from deep states of quiescence may appear surprising because ASCL1 protein is typically only detected in active NSCs when using conventional antibody staining techniques. However, despite being broadly transcribed, ASCL1 is typically only detected in active NSCs because it undergoes proteasomal degradation due to destabilisation by inhibitor of DNA binding (ID) transcription factor proteins in quiescent NSCs^11^. Despite these post-translational mechanisms controlling ASCL1 levels in quiescent NSCs, it is clear that small amounts of protein are produced. For example, using sensitive antibodies against a VENUS fusion protein we have previously detected ASCL1 protein in more than 50% of quiescent NSCs, at least in young mice^11^.

Conversely, we found that the *Mycn* transcription factor promotes activation from shallow states of quiescence. *Mycn* appears to be important in increasing biosynthesis, consistent with its role in other systems such as in embryonic diapause^20^ or in quiescent haematopoietic stem cells^19^. We analysed conditional genetic knockout mice of *Mycn, Myc* and *Mycl* side-by-side and only observed significant defects in *Mycn* mice, and the phenotype of triple knockout mice was comparable to loss of *Mycn* alone. While we cannot rule out minor roles for *Myc/Mycl* in the activation process, indeed *Myc* has recently been suggested to have a role in promoting hippocampal NSC activation^31^, clearly *Mycn* is the dominant family member in this context. These sequential transcriptional programs governing progression from deep quiescence to shallow quiescence are likely linked through direct transcriptional regulation, where ASCL1 activates the expression of MYCN. ASCL1 has previously been shown to transcriptionally regulate MYCN in other scenarios. For example, MYCN is a target gene of ASCL1 in glioblastoma^32^. Similarly, in neuroblastoma, ASCL1 acts as a pioneer factor where it then recruits and cooperates with MYCN at genomic loci to drive adrenergic cell fate conversion^33^. Whether ASCL1 and MYCN also drive gene expression in a cooperative manner in quiescent NSCs is unclear, although relatively few genes were commonly mis-regulated between both datasets, which would argue for limited cooperation. Regardless of the exact functional relationship between these transcription factor proteins, a useful mRNA combinatorial code can be generated to classify hippocampal NSCs into deep states of quiescence (*Ascl1+Mycn-Ki67-*), shallow states of quiescence (*Ascl1+Mycn+Ki67-*) and active states (*Ki67+*). Future studies should explore whether *Ascl1*+*Mycn+Ki67–* hippocampal NSCs encompass all cells in shallow quiescence or whether populations such as resting NSCs^6^ and primed NSCs^4^ represent distinct populations.

### Limitations of study

Interestingly, MYCN also serves as a substrate for HUWE1-mediated ubiquitination during embryonic brain development^34^. While our prior work established ASCL1 as the primary target of HUWE1 in hippocampal NSCs^10^, the possibility that elevated MYCN protein levels contribute to the phenotype observed in HUWE1 knockout mice cannot be entirely ruled out. However, our conditional deletion of the *Mycn* gene demonstrates that the primary function of *Mycn* is to promote progression through shallow quiescence.

## Acknowledgments

We gratefully acknowledge Lan Chen and the Biological Resources, Advanced Sequencing, Flow Cytometry, and Light Microscopy Facilities; and the Research Illustration team of the Francis Crick Institute for technical support. This work was supported by the Francis Crick Institute, which receives its funding from Cancer Research UK (FC0010089), the UK Medical Research Council (FC0010089), and the Wellcome Trust (FC0010089). This work was also supported by QIMR Berghofer.

## Author contributions

Conceptualization, F.G and LH; Data Curation, L.H.; Formal Analysis, L.H., P.R., S.A.P., R.L.J., C.C; Investigation, L.H., P.R., S.A.P., R.L.J., Visualization, L.H., P.R., S.A.P., R.L.J., Methodology, L.H., R.L.J., F.G.; Writing – Original Draft, L.H.; Writing – Review & Editing, all authors; Supervision, F.G. and L.H.; Funding Acquisition, F.G. and L.H.; Project Administration, L.H.

## MATERIALS AND METHODS

### Lead contact

Further information and requests for reagents should be directed to and will be fulfilled by the Lead Contact, Lachlan Harris.

### Materials availability

This study did not generate new reagents.

### Data and code availability

scRNA-seq sequencing files (FASTQs), cellranger outputs and original code will be made available upon publication.

## EXPERIMENTAL MODEL AND SUBJECT DETAILS

All mice were maintained on a mixed genetic background. The Ascl1 cKO^9^, Myc cKO^21^, MycN cKO^22^, MycL cKO^23^; Nestin-GFP^36^, YFP^12^ and Glast-creERT2^13^ transgenic lines have all been previously reported. Both male and female mice were used throughout the study, an exception to this was for the x-linked Huwe1 conditional allele, where only males were used. The effects of the gene knockouts seen throughout the study were consistent across sexes. All experimental protocols involving mice were performed in accordance with guidelines of the Francis Crick Institute, national guidelines and laws. This study was approved by the UK Home Office (PPL PB04755CC). Throughout the study, mice were housed in standard cages with a 12 h light/dark cycle and ad libitum access to food and water.

## METHOD DETAILS

### Preparation and sectioning of mouse brain tissue

Mice were perfused transcardially with phosphate buffered saline (PBS), followed by 4% paraformaldehyde (10-20 mL) and postfixed for 16-24 h before long term storage in PBS with 0.02% sodium azide, at 4°C. Brains were sectioned in a coronal plane at 40 µm using a vibratome (Leica). The entire rostral-caudal extent of the hippocampus was collected in a 1 in 12 series.

### Antibodies, immunofluorescence, cell counts

At minimum, 1 series per mouse (5-6 sections) was stained and analysed per experiment. Free-floating sections underwent antigen retrieval in sodium citrate solution (10mM, pH6.0) for 10min at 95°C. The retrieval time was reduced if the detection of endogenous YFP/GFP was required, as these proteins are heat sensitive and cannot be re-detected with commercially available antibodies against GFP ^37^. Sections were blocked with normal donkey serum (2%) diluted in PBS-Triton X-100 (0.2%) for a minimum of 2 h. Sections were then incubated overnight at 4°C with primary antibodies diluted in blocking buffer. After three washes in PBS for 10 min, the sections were incubated with fluorescent secondary antibodies at room temperature for 2 h (Jackson ImmunoResearch). The sections were counterstained with 4’,6- diamidino-2-phenylindole (DAPI, ThermoFisher Scientific) and mounted onto Superfrost slides.

The sections were imaged using a SP5/SP8 Leica confocal microscope with a 40X objective or a Olympus CSU-W1 SoRa spinning disc confocal using a 20X objective. Cell counts were then performed, data was normalised to number of cells per surface area of subgranular zone, which was measured by the length of the subgranular zone in coronal brain sections multiplied by the thickness of the vibratome section. In all analyses, we identified NSCs as those cells that had a radial GFAP-positive process that we could confidently link to a SOX2-positive nucleus in the subgranular zone (SGZ). The cells were classified as quiescent or proliferating depending on the expression of Ki67. To calculate the proportion of NSCs expressing MYCN in control mice at minimum 20 quiescent and proliferating NSCs were examined per animal. To calculate the proportion of NSCs expressing MYCN in *Ascl1* cKO versus *Ascl1* control mice a minimum of 40 cells were examined per animal. Cell counts were not performed blinded to genotype, as the severe phenotypes of *Ascl1*, *Huwe1*, *Mycn*, and triple *Myc* cKO mice made blinding of images impractical.

### Tamoxifen treatment

Tamoxifen solution (10-20 mg/mL) was prepared for oral gavage by dissolving the powder (Sigma-Aldrich) in a mix of 10% ethanol and 90% cornflower oil (Sigma-Aldrich) and was provided to mice via oral gavage (100 mg/kg).

### Single-cell RNA sequencing

#### Sample Preparation

Each independent scRNA-seq experiment (batch) comprised a pooled group of 1-2 conditional knockout mice (of either *Ascl1*, *Huwe1* or *Mycn cKO* genotype) with 1-2 littermate controls that were prepared simultaneously to minimise processing artefacts. In total, 3 independent experiments were performed on *Ascl1* cKO mice and controls, 2 independent experiments on *Huwe1* cKO mice and controls and 2 independent experiments on *Mycn* cKO mice and controls (Table S1). In the *Ascl1* experiments, the cKO mice were homozygous for the floxed allele, heterozygous for Glast-creERT2 and homozygous for the cre-reporter YFP allele, while the the control mice were genetically identical but were wildtype for *Ascl1* alleles. In the *Huwe1* experiments, the male cKO mice were hemizygous for the floxed allele, heterozygous for Glast-creERT2 and homozygous for the cre-reporter YFP allele, while the male control mice were genetically identical but were wildtype for *Huwe1*. All mice received tamoxifen via oral gavage (100mg/kg) for 5 days starting from P30 (range from P27-P33) and were euthanised 12-days later. For the *Mycn* experiments, the cKO and control mice were homozygous for the floxed allele, heterozygous for Glast-creERT2 and heterozygous for Nestin-GFP. The *Mycn* cKO mice received tamoxifen and *Mycn* control mice received corn-oil via oral gavage for 5- days at P30 (range from P27-P30) before being euthanised 12-days later. In the majority of experiments, at least two cKO or control mice were combined to guarantee the sorting and sequencing of sufficient numbers of cells (Table S1).

Mice were euthanised by cervical dislocation, and the hippocampal dentate gyri were dissected^38,39^. The dentate gyrus was disassociated using the Neural Tissue dissociation kit (P) (Milteny Biotec) with the following exceptions, a 37C orbital shaker was used during the enzymatic digestions, and we used manual trituration with fire-polished pipettes to aid dissociation following the incubations with enzymatic mix 1 and 2^6^. Cells were also live-dead counterstained with DAPI. The cells were then sorted on a MoFlo XDP (Bechman Coulter) using a 100 µm nozzle. Debris were removed, followed by two gates to remove aggregates and dead cells, based on DAPI fluorescence. Cells were then gated for YFP or GFP expression according to cells from a control mouse hippocampus that did not contain a fluorescent transgene. We sorted between 10,000-25,000 cells per group (2 mice per group) over a period of 15-30 min per sample. The cells were sorted into 700 µl of recovery media (0.5% PBS-BSA in DMEM/F-12 without phenol red) in 1.5mL tubes, and spun down at 500 *g* for 7 min at 4°C. The cells were then gently resuspended in the residual recovery media using a wide-bore pipette to a final volume of 50 µl. The single-cell suspension (to a maximum of 10,000 cells) was then loaded into the 10x Chromium.

#### Sequencing and data processing

On each experimental day, two libraries were prepared, one for each of the experimental groups to control for batch effects, which are strong in this type of data (i.e., cKO and control). All libraries were prepared with 10x Genomics Chemistry, Single Cell version 3.0.1. After sequencing, cellranger count (Ver. 7.0.1) was used to map the FASTQ files from the Ascl1 and Huwe1 experiment to refdata-gex-mm10-2020-A. In contrast, the Mycn data was mapped to a custom version of this genome, which had the *Mycn* gene deleted and replaced with two new contigs, corresponding to exon1 of the *Mycn* gene (*Mycnex1*) and exon 2/3 (*Mycnex3*). This modification was made to detect cells that had undergone recombination of the *Mycn* locus. Specifically, in control cells, the majority of reads map to the exon 2 and 3 of the *Mycn* locus, which are directly upstream of the polyA tail. In contrast, in cells that have recombined *Mycn*, exons 2 and 3 are deleted, and all reads map to exon 1. The gene encoding eGFP was also added to the custom genome. The genome was made with the cellranger mkref command. As both males and female mice were used in this study but sex differences were not the focus of our investigation, the Y chromosome genes with highest expression (*Eif2s3y*, *Ddx3y*, *Gm47283*, *Uty* and *Kdm5d*) and genes involved in X-inactivation (*Xist* and *Tsix*) were deleted from the count matrices prior to normalisation and clustering.

#### Quality control

Individual samples were read into R version 4.2.0 and analysed using Seurat (Ver. 4.3.0.1)^40,41^. Unless otherwise specified, plots were created using ggplot2 version 3.4.2. GEM beads were excluded if they nCount_RNA values were less than 600-5000 (varied according to distribution of data within individual samples) or had greater than 8% mitochondrial reads (uniform across all samples). In total, 77.9% of cells passed quality control in Ascl1 data, 78.8% in Huwe1 data, and 94.9% in Mycn data. In the second *Ascl1* cKO replicate (comprising 1 male, 1 female) we found that a female mouse from the control group (compromising 1 male, 1 female) had been inadvertently swapped with the female *Ascl1* cKO mouse. Cells from the *Ascl1* cKO and control samples were reassigned to their correct experimental groups by first identifying the individual mouse each cell originated from, based on sex-specific gene expression. To do this, a sex score was calculated by subtracting raw counts of the sum of female genes involved in X-inactivation (*Xist* and *Tsix*) from Y chromosome genes (*Ddx3y*, *Uty* and *Eif2s3y*). A negative sex score indicated female sex and a score of ≥ 0 indicated male sex. Subsequently, the cells were assigned to the appropriate genotype by confirming the presence or absence of *Ascl1* expression.

#### Normalisation, dimensionality reduction and clustering

All *Ascl1* cKO and control samples were merged into one large Seurat object. Separately, this was repeated for all *Huwe1* samples and all *Mycn* samples. The merged Seurat objects were then split according to experimental replicate using the function SplitObject and the data transformed using the SCTransform function, with the vst.flavor argument set to “v2”. The data was integrated using the function SelectIntegrationFeatures with the nFeatures argument set to “3000” and the functions PrepSCTIntegration, FindIntegrationAnchors and IntegrateData using default parameters to control for batch effects between experimental days. Elbow plots were used to approximate the number of principal components for clustering and visualisation using UMAP plots^42^. As axis values of UMAPs are not meaningful, we orientated the final plots so that active NSC clusters appear towards the top-right of each plot and/or differentiation processes proceed from left-to-right.

After identification of clusters, we isolated the three neurogenic clusters according to known marker expression, quiescent NSCs (*Hopx*-high, *S100b*-low), proliferating cells (containing cell-cycle markers, NSC markers and intermediate progenitor cell markers), and neuroblasts (Dcx-high, Eomes-low). After isolating these cells, we used the SCTransform workflow to recluster these populations. Next, we sought to isolate only NSCs. As previously described, the proliferating cell cluster contained both proliferating NSCs and intermediate progenitor cells^6^. Therefore, we next separated proliferating NSCs from intermediate progenitor cells based on marker expression (*Hopx/Apoe-*high, *Dcx/Neurod2-*negative; *Eomes/Tubb3*-low) and reiterated the SCTransform workflow. We repeated this process by removing proliferating NSCs and reclustered a final time. The result was four Seurat objects (all cells, neurogenic clusters, NSCs and quiescent NSCs) per *Ascl1*, *Huwe1* and *Mycn* data set. For downstream trajectory inference and pseudobulk analysis, Seurat objects were converted to SingleCellExperiment objects using the Seurat function as.SingleCellExperiment with the assay argument set to “RNA”.

#### Pseudo-bulk differential expression analysis

A pseudo-bulk approach was used to perform differential gene expression analysis between cKO and control conditions. Pseudo-bulk samples were created by aggregating counts across individual samples using the scuttle (version 1.8.4) function aggregateAcrossCells. Differential expression analysis was performed using the scran (version 1.26.2) function pseudoBulkDGE, which is a wrapper for edgeR’s quasi-likelihood methods. To run pseudoBulkDGE, the “edgeR” method was used, and batch effects between biological replicates (experiments) were accounted for using the design formula “∼Batch + Genotype”, where Batch represented the biological replicate, and Genotype represented cKO or control. Differentially expressed genes were determined using false discovery rate (FDR) < 0.05. Transcription Factor (TF) status (0 and 1) of each gene was determined based on a previously published classification^43^.

#### Gene Ontology enrichment analysis

Gene Ontology (GO) enrichment analysis was performed using clusterProfiler (version 4.6.2)^44^. First, gene symbols for differentially expressed genes identified by pseudo-bulk analysis were converted to Entrez IDs using the bitr function. The Entrez IDs were then passed to the enrichGO function with ont argument set to “BP” (Biological Process category). Significant terms were determined using FDR < 0.05.

#### Pseudotime analysis

For pseudotime analysis the quiescent NSC Seurat objects were used. The quiescent NSCs were ordered from deep to shallow quiescence using the pseudotime inference tool Slingshot version 2.6.0^17^. Specifically, the slingshot function was used with reducedDim argument set to PCA. To ensure the orientation of the trajectory was correct, we assessed the expression of quiescence marker genes (*Apoe*, *Clu*, Hopx, *Id4* and *Sparc*) and activation marker genes (*Ccnd1*, *Ccnd2*, *Rpl10* and *Thrsp*), where we expect increased expression of the markers at the start and end of the trajectory, respectively. Whilst Slingshot automatically placed the trajectory for the *Mycn* samples in the correct orientation, the trajectory for the *Ascl1* samples was in the reverse orientation. To overcome this, we created manual cluster labels based on Seurat clusters, where all but the Seurat cluster that should be the end cluster was labelled as one cluster. These manual cluster annotations were then used for the clusterLabels argument to run slingshot.

#### Trajectory-based differential expression analysis

Downstream of trajectory inference, we performed trajectory-based differential expression analysis between experimental groups (cKO versus control) using tradeSeq version 1.12.0^25,26^. First, we evaluated the number of knots to use for the negative binomial generalised additive models (NB-GAMs) with the evaluateK function using default settings. Based on the graphical output from evaluateK, we purposely chose the same number of knots (6 knots) across the *Ascl1* and *Mycn* experiments to enable comparisons. Next, we ran the fitGAM function to fit NB-GAMs to the SingleCellExperiment data, with nknots = 6 and conditions argument specifying Genotype (cKO and control). Knots were visualised in reduced dimensions (PCA) using plotGeneCount. We then ran the conditionTest function multiple times to test for differential expression between experimental groups along pseudotime. For conditionTest, we used l2fc value of log2(1.2), as we found hundreds to thousands of significant genes without such a threshold, and for each test we contrasted sequential knots using the knots argument (vector of two), namely knots 1 vs 2, 2 vs 3, 3 vs 4, 4 vs 5, 5 vs 6. Results were sorted by descending Wald statistic, and p-values were adjusted using p.adjust “fdr” method. Significant genes were determined using FDR < 0.05, and visualized using plotSmoothers.

#### ChIP-sequencing analysis

The ASCL1 ChIPseq fastq file (ERR376162) was obtained from Andersen and colleagues ^9^ (deposited in the European Nucleotide Archive’s Sequence Read Archive under accession number ERP004380), the p300 (ERR216112) and H3K27ac (ERR216108) ChIPseq fastq files were obtained from Martynoga and colleagues^35^ (deposited in the European Nucleotide Archive’s Sequence Read Archive under accession number ERP002084).

Raw reads data were quality-controlled using FastQC (v. 0.11.8). Reads were aligned using Bowtie2 (v. 2.5.1) to the mm10 mouse reference genome using default parameters. Samtools (v.1.18) was then used to convert sam files to bam format. Sambamba (v. 0.6.6) was used to sort bam files by genomic coordinates and filter out PCR duplicates and unmapped reads to contain only uniquely mapping reads. deepTools (v. 3.5.1) was used to obtain the bigWig files.

## QUANTIFICATION AND STATISTICAL ANALYSIS

### Statistical analysis of cell counts

The statistical testing approach were implemented using Graphpad Prism (version 8.0) or in R (4.2.0). Two-tailed unpaired Student’s t tests were performed when comparing two groups. For experiments involving two independent variables, a two-way ANOVA was performed. Any significant main effect detected by ANOVA was followed by multiple comparisons analysis using multiple t-test and a pooled estimate of variance where appropriate with Holm-Sidak correction.

### Single-cell data

To determine if the distribution of quiescent NSCs along pseudotime were different between experimental groups, a Kolmogorov-Smirnov test was performed with function ks.test in R (4.2.0).

### List of antibodies

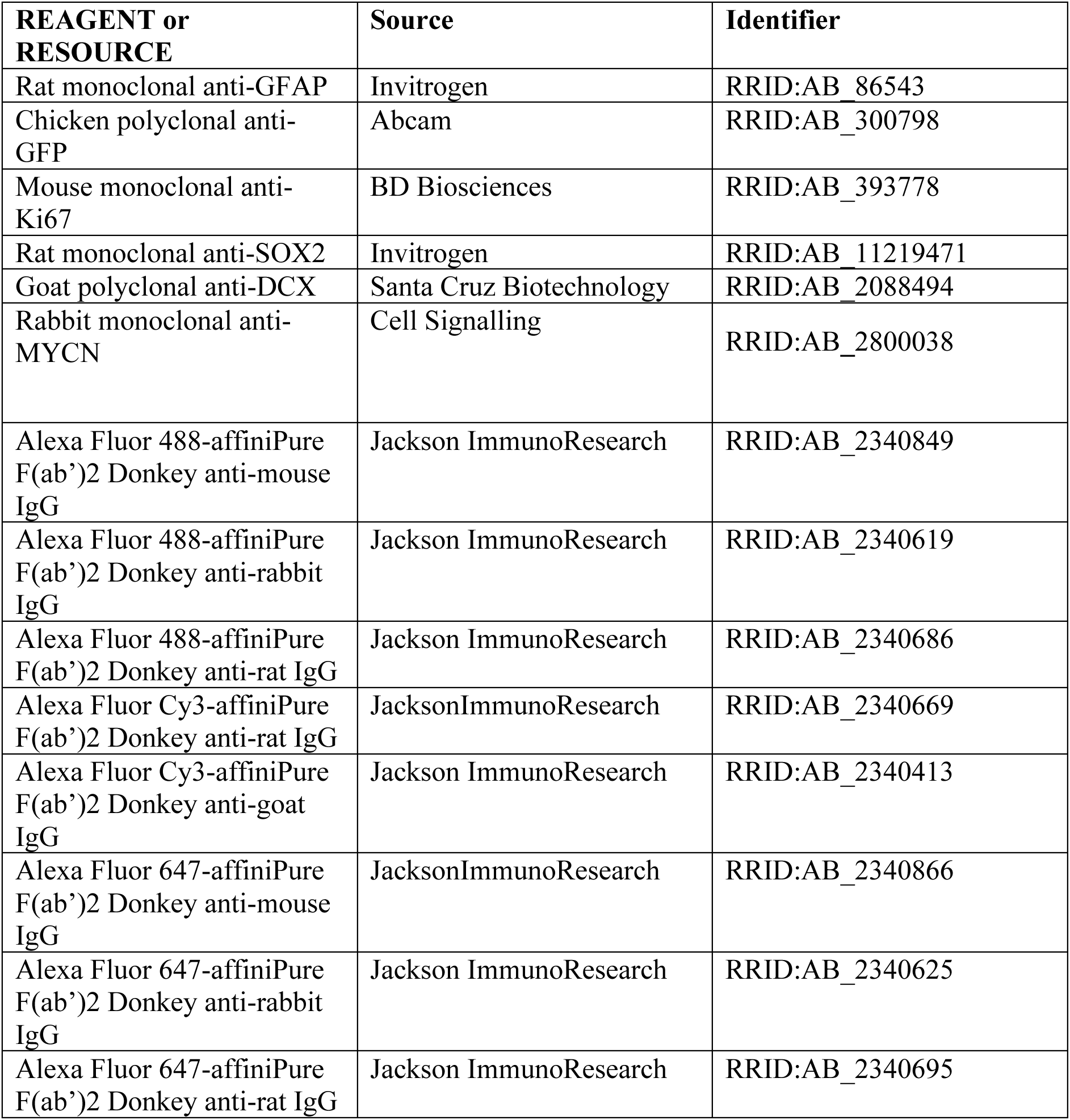

## SUPPLEMENTAL ITEMS

**Figure S1:**
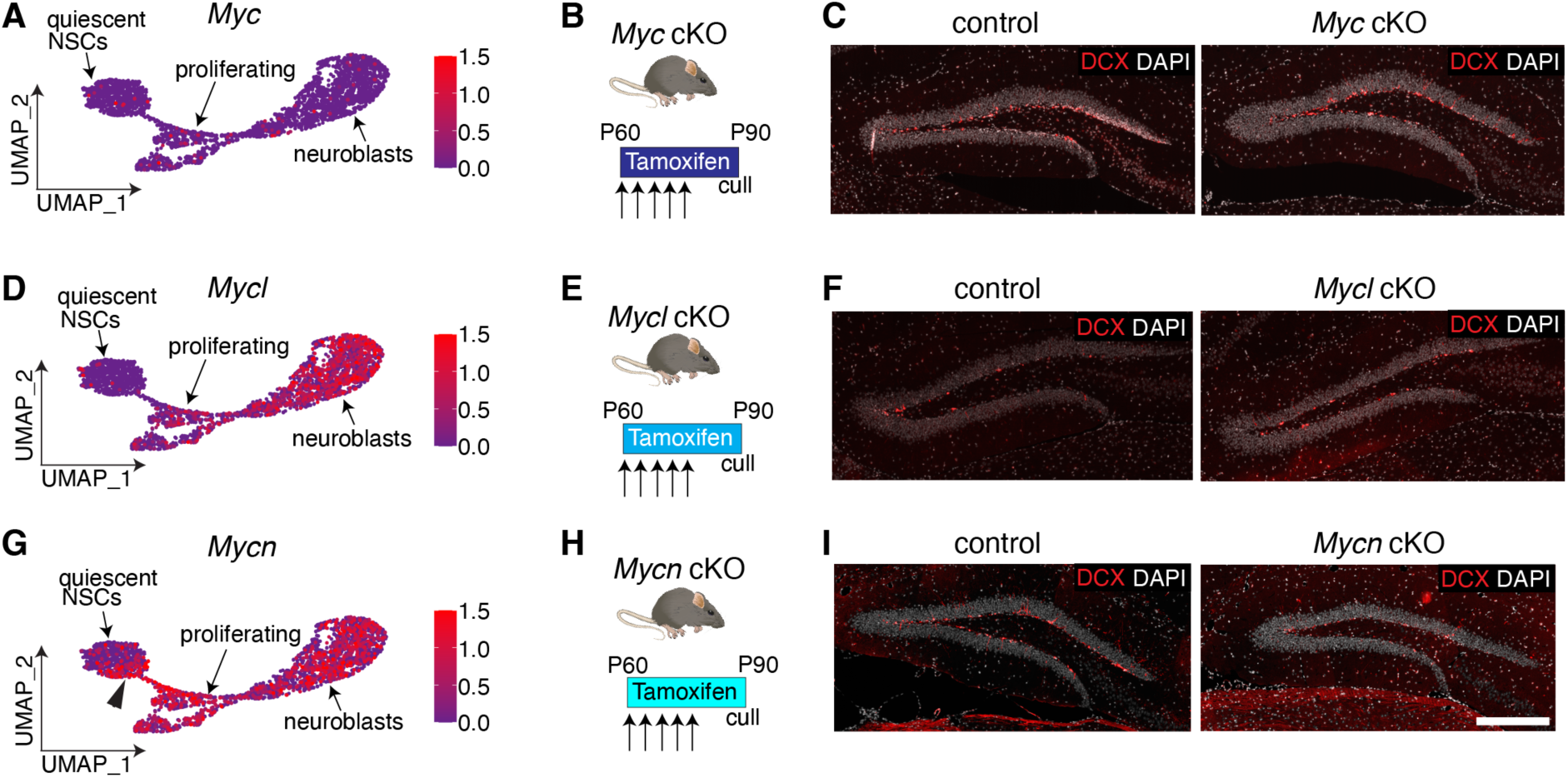
Loss of *Myc* or *Mycl* does not substantially impair adult hippocampal neurogenesis, related to Figure 3. (A) Expression of *Myc* in adult hippocampal neurogenic lineage, data from *Ascl1* control mice in Figure 1. (B) *Myc* cKO and control mice were injected with tamoxifen for 5 days and culled 30 days later. (C) DCX staining in *Myc* cKO mice was comparable to controls. (D) Expression of *Mycl* in adult hippocampal neurogenic lineage, data from *Ascl1* control mice in Figure 1. (E) *Mycl* cKO and control mice were injected with tamoxifen for 5 days and culled 30 days later. (F) DCX staining in *Mycl* cKO mice was comparable to controls. (G) Expression of *Mycn* in adult hippocampal neurogenic lineage, data from *Ascl1* control mice in Figure 1. (H) *Mycn* cKO and control mice were injected with tamoxifen for 5 days and culled 30 days later. (I) DCX staining in *Mycn* cKO mice was substantially reduced compared to controls. Scale bar (located in I): 320 μm in (C, F, I). Arrowhead (located in G) indicates group of quiescent NSCs expressing *Mycn* but not *Myc* or *Mycl*.

**Figure S2:**
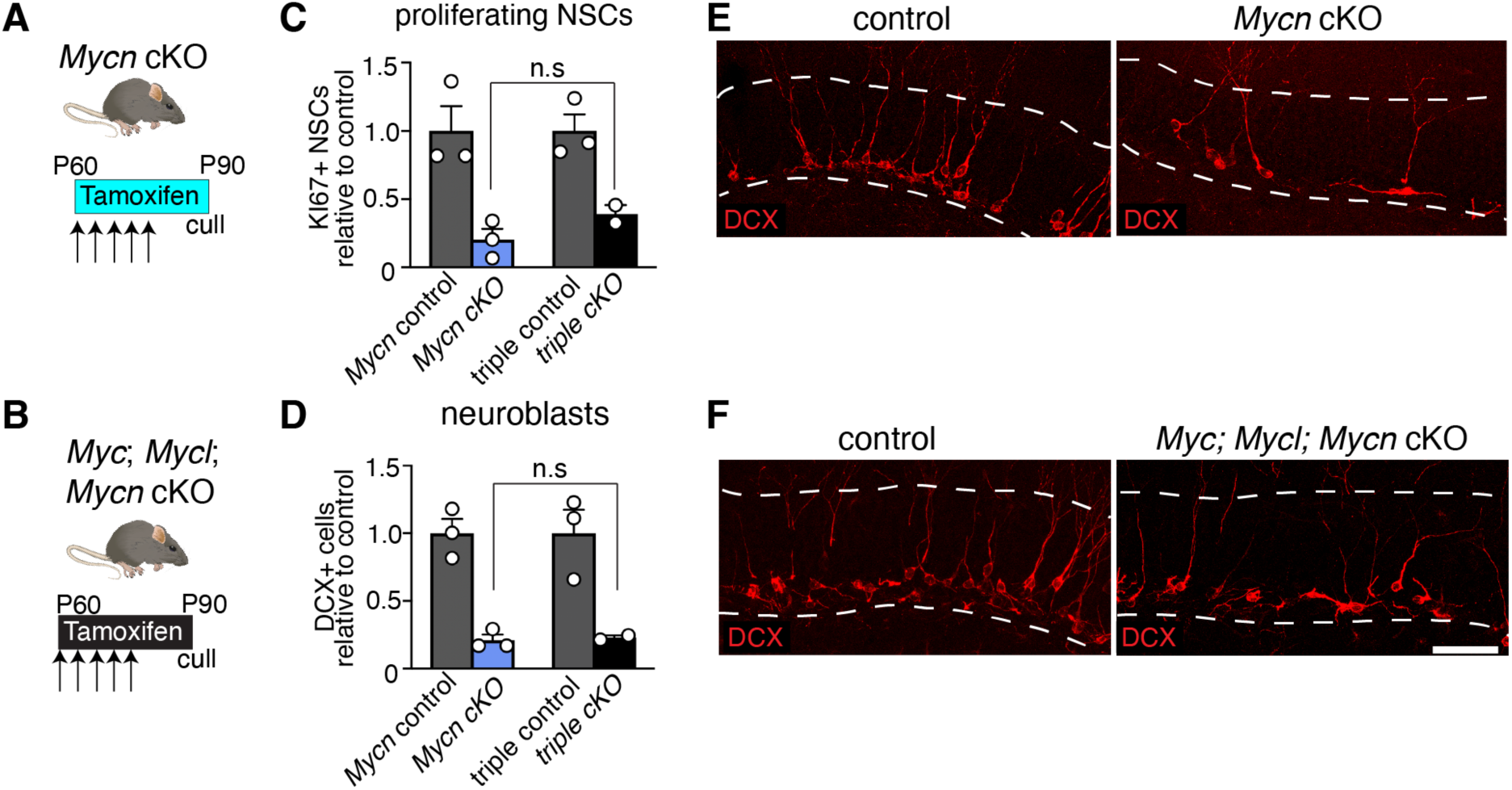
Loss of three *Myc* genes from adult neural stem cells is comparable to loss of *Mycn* alone, related to Figure 3. (A) *Mycn* cKO and control mice were injected with tamoxifen for 5 days and culled 30 days later. (B) Triple *Myc* cKO and control mice were injected with tamoxifen for 5 days and culled 30 days later. (C) Fold change in proliferating NSC number in *Mycn* cKO and triple *Myc* cKO mice relative to respective controls. (D) Fold change in neuroblast number in *Mycn* cKO and triple *Myc* cKO mice relative to respective controls. (E) DCX staining in *Mycn* cKO and control mice. (F) DCX staining in triple *Myc* cKO and control mice. Graphs in C, D show mean +/- SEM. Statistics: t-test (C, D). Scale bar (located in F): 60 μm in (E, F). Dashed lines demarcate granule cell layer in (E, F).

**Figure S3:**
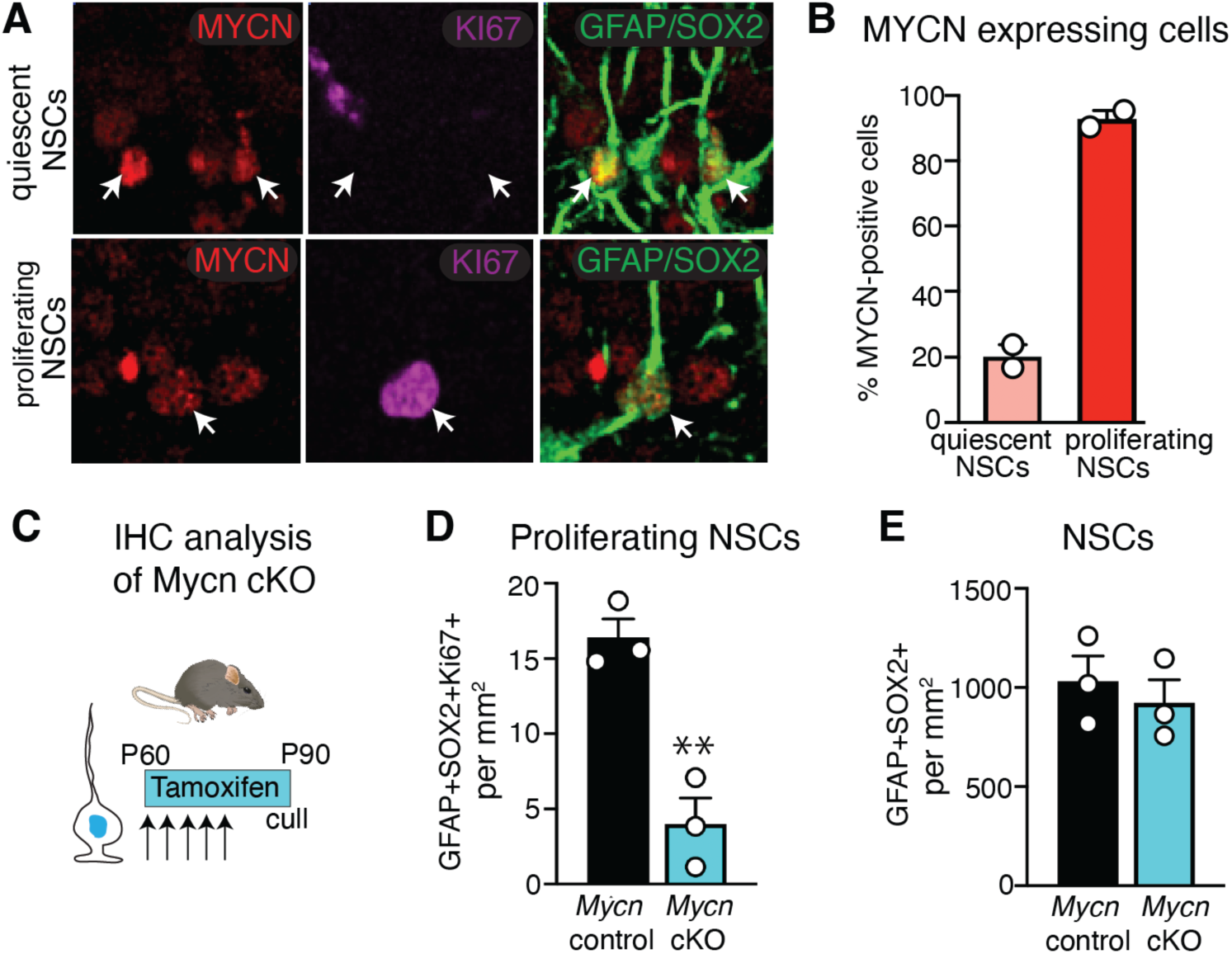
*Mycn* loss reduces the number of proliferating NSCs, related to Figure 3. (A) MYCN protein is detected in quiescent NSCs (upper panels) and proliferating NSCs (lower panels) in the adult hippocampus. (B) Quantification of the proportion of quiescent and proliferaring NSCs expressing MYCN by cell-type. (C) Conditional, inducible deletion of *Mycn* (Glast-creERT2; *Nestin::GFP*) followed by immunofluorescence analysis. *Mycn* cKO and control mice received tamoxifen for 5 days and were culled 30 days later. (D) Quantification of the number of proliferating NSCs in *Mycn* cKO mice and controls. (E) Quantification of the number of NSCs in *Mycn* cKO mice and controls. Graphs in D, E show mean +/- SEM. Statistics: t-test (D, E). \*\**P* < 0.01.

**Figure S4:**
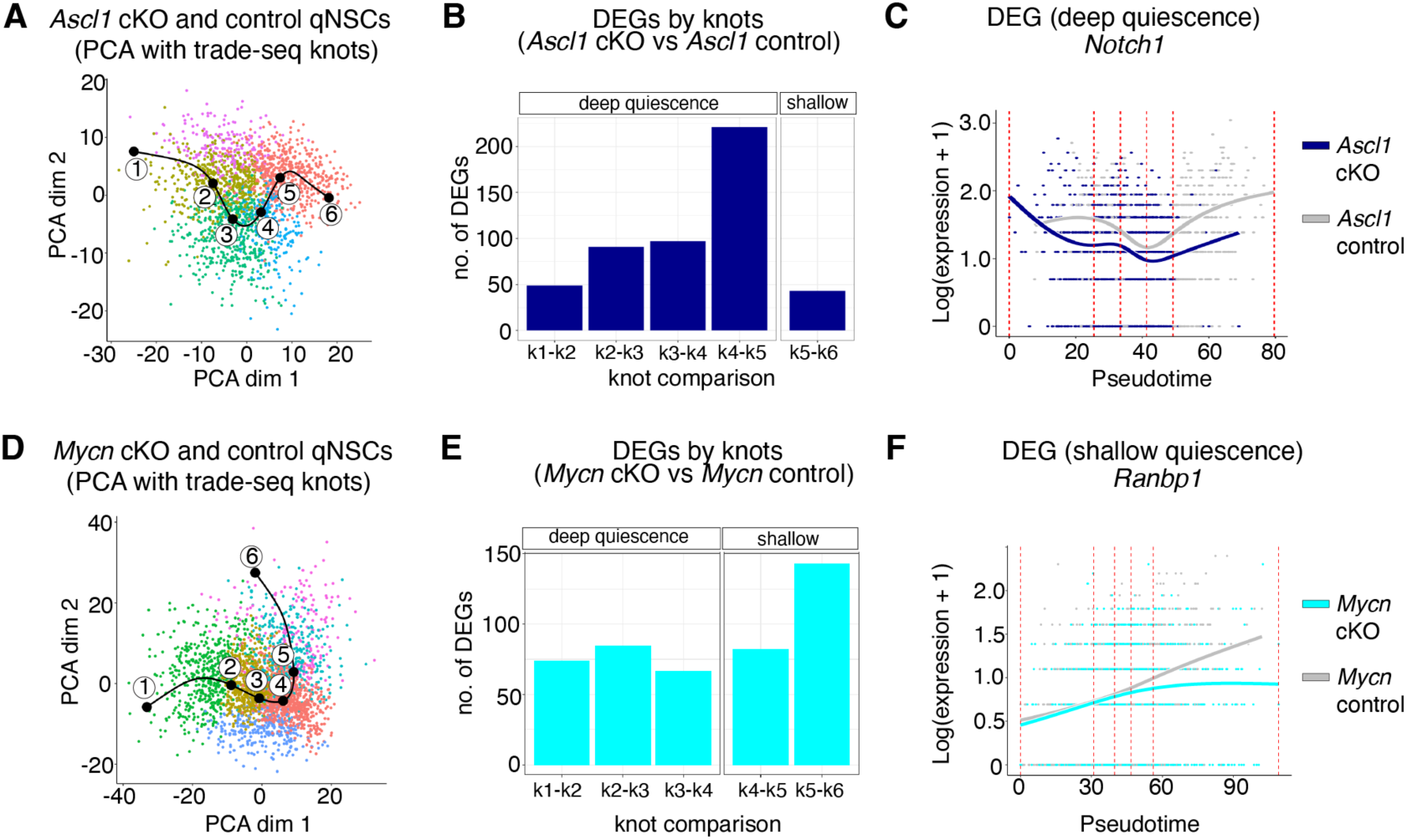
*Ascl1* loss impacts deeper stages of NSC quiescence than *Mycn* loss, related to Figure 4. (A) Principal Component Analysis (PCA) dimensions 1 and 2 of *Ascl1* cKO and control quiescent NSCs, labelled with six knots marking binning of cells into five equally sized partitions of pseudotime as determined by tradeSeq. Cells coloured by Seurat cluster. (B) Number of differentially expressed genes (DEGs) at each knot comparison between *Ascl1* cKO and control quiescent NSCs. Deep quiescence and shallow quiescence defined as knots that occur before and after median pseudotime position of control cells, respectively. (C) Example of a DEG from early knot comparisons (deep quiescence) between *Ascl1* cKO and control quiescent NSCs. (D) PCA dimensions 1 and 2 of *Mycn* cKO and control quiescent NSCs, labelled with six knots marking binning of cells into five equally sized partitions as determined by tradeSeq. Cells coloured by Seurat clusters. (E) Number of DEGs at each knot comparison in *Mycn* cKO and control quiescent NSCs. Deep quiescence and shallow quiescence defined as knots that occur before and after median pseudotime position of *Mycn* cells, respectively. (F) Example of a differentially expressed gene from late knot comparisons (shallow quiescence) between *Mycn* cKO and control quiescent NSCs

